# Conserved residues in the extracellular loop 2 regulate *Stachel*-mediated activation of ADGRG2

**DOI:** 10.1101/2021.02.14.431159

**Authors:** Abanoub A. Gad, Pedram Azimzadeh, Nariman Balenga

## Abstract

Cleavage and dissociation of a large N-terminal fragment and the consequent unmasking of a short sequence (*Stachel*) remaining on the N-terminus have been proposed as mechanisms of activation of some members of the adhesion G protein-coupled receptor (aGPCR) family. However, the identity of residues that play a role in the activation of aGPCRs by the cognate *Stachel* remains largely unknown. Protein sequence alignments revealed a conserved stretch of residues in the extracellular loop 2 (ECL2) of all 33 members of the aGPCR family. ADGRG2, an orphan aGPCR, plays a major role in male fertility, Ewing sarcoma cell proliferation, and parathyroid cell function. We used ADGRG2 as a model aGPCR and generated mutants of the conserved residues in the ECL2 via site-directed mutagenesis. We show that tryptophan and isoleucine in the ECL2 are required for receptor stability and surface expression in the HEK293 cells. By adjusting the receptor surface expression levels, we show that the ECL2 mutation ablates the *Stachel*-mediated activation of multiple signaling pathways of ADGRG2. This study provides a novel understanding of the role of the ECL2 in *Stachel*-mediated signaling and degradation of ADGRG2, which may lay the foundation for the rational design of therapeutics to target aGPCRs.

## Introduction

G protein-coupled receptors (GPCRs) play fundamental roles in various cellular processes such as proliferation, metabolism, hormone secretion, contraction, and neurotransmission^1^. The signaling pathways of GPCRs via G proteins and β-arrestins, and their trafficking trajectories have been studied for several decades^2,3^. This in combination with recent advances in our understanding of GPCR structures has formed a strong foundation for rational drug design to target this largest superfamily of surface receptors^4,5^.

Recent genomic and model organism studies have revealed the versatile roles that the adhesion GPCRs (aGPCRs), the second-largest family of GPCRs, play in endocrine, nervous and immune systems^6^. aGPCRs have structural differences with other classes of GPCRs, namely autoproteolysis at a GPCR proteolysis site (GPS) and formation of a two-segmented receptor with a large N-terminal fragment (NTF) and a C-terminal fragment (CTF). The CTF includes the C-terminus, seven-transmembrane helical domains (TM1-7), extracellular loops (ECL1-3), intracellular loops (ICL1-3), and a short extracellular sequence on the N-terminus just before the TM1. Many studies^7–10^, including our recent reports^11,12^, have shown that this short remaining peptide, known as *Stachel* sequence, can activate aGPCRs and might be the mechanism of activation in physiological and pathological states. NTF-dissociating molecular partners can potentially unmask the *Stachel* sequence for its binding to the cognate aGPCR. These studies exploited two main tools to reveal this mechanism: (a) NTF-truncated aGPCRs that show constitutive activation of G proteins; (b) synthetic peptides resembling the *Stachel* sequence that activate aGPCRs^13,14^.

We recently showed that human ADGRG2 (GPR64), an orphan aGPCR, is expressed in human parathyroid glands and regulates the signaling and function of calcium-sensing receptor^12^. We discovered that similar to other aGPCRs (ADGRG1^15^, ADGRD1^8^, ADGRG6^8^), ADGRG2 is activated by either the endogenous 15-amino acid long *Stachel* (P-15) on the N-terminus of its NTF-truncated mutant (ADGRG2-ΔNTF) or the synthetic P-15^12^. We showed that the deletion of P-15, in addition to the NTF, ablates constitutive activation of Gαs and cAMP production in a mutant that starts with the proline 622 (ADGRG2-P622)^11^ and elevates receptor response to synthetic P-15. Nevertheless, the binding site of *Stachel* remains unknown among aGPCRs.

Previous studies in other families of GPCRs revealed that some residues in the ECL2 play major roles in ligand access, receptor subtype selectivity, and activity^16–18^. Despite high degrees of diversity in the structure of ECL2 among GPCRs^19^, there is a conserved disulfide bond between the cysteines of ECL2 and TM3, which ensures receptor structural integrity. In the Secretin family (class B1), this conserved ECL2 cysteine is followed by a tryptophan residue, forming the CW motif, which is further followed by an acidic residue (aspartic or glutamic acid). The aGPCR family has the highest homology to the Secretin family. By aligning the ECL2 residues of all 33 human aGPCRs (predicted by either GPCRdb or Uniprot), we show that most of the aGPCRs have an aliphatic residue (leucine or isoleucine) after the CW motif (Figure 1a and Supplementary Figure 1). Whether the ECL2 plays a role in the activation of aGPCRs by *Stachel* remains unknown.

**Figure 1.**
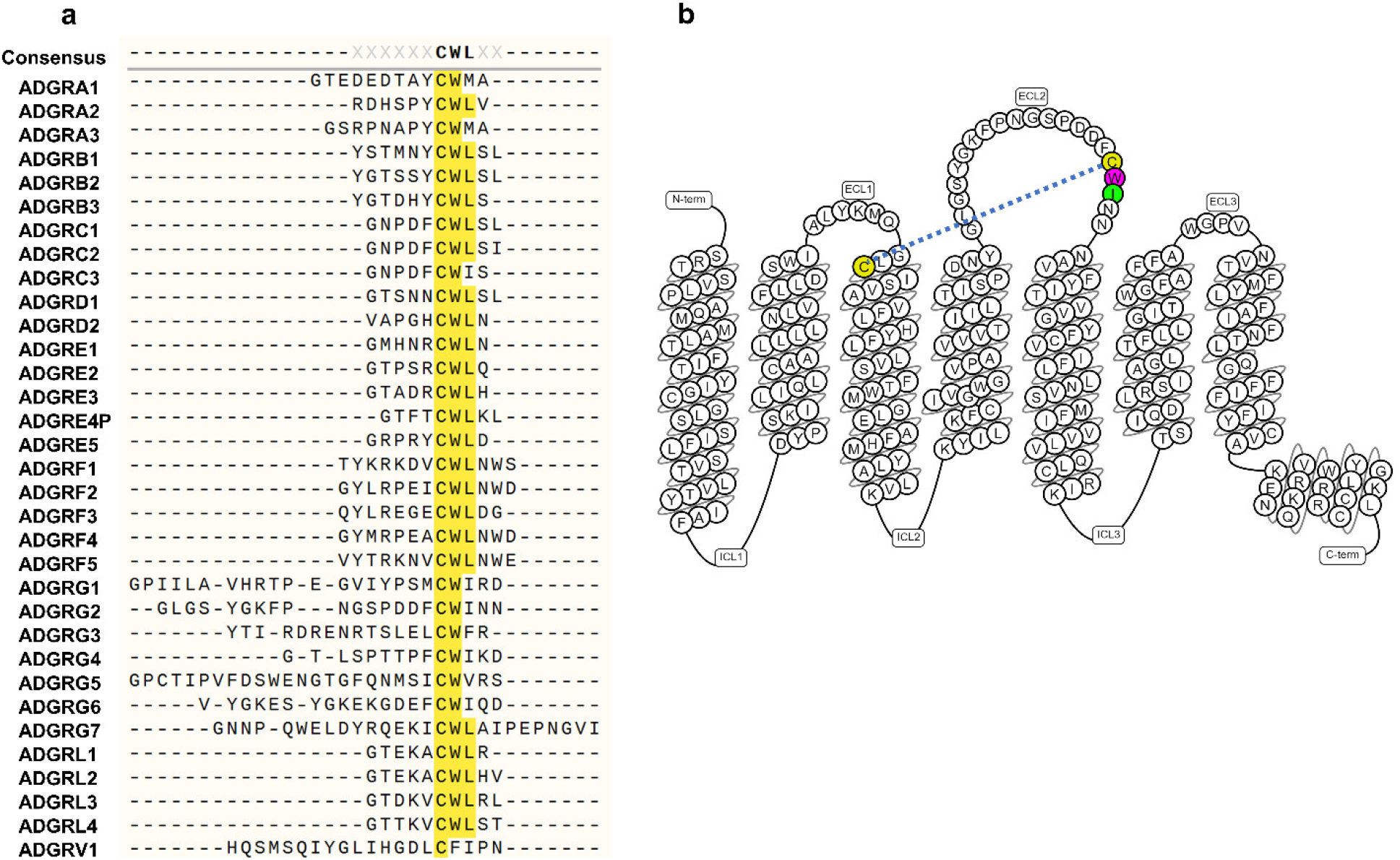
Conserved residues in the extracellular loop 2 (ECL2) of aGPCRs. (a) Multiple alignments of the ECL2 of all 33 members of the aGPCR family. Predicted amino acid sequences of the ECL2 for each aGPCR were derived from GPCRdb. The alignment was conducted in SnapGene software using the Clustal Omega algorithm^42^. (b) Snakeplot of human ADGRG2, exported from GPCRdb, showing the colored CWI motif in the ECL2. The sequences of the N-terminal fragment and C-terminus are not shown. The dashed line shows the disulfide bond between the cysteines of TM3 and ECL2.

Here, we use ADGRG2-P622 as a model aGPCR to investigate the role of tryptophan and isoleucine of ECL2 (Figure 1b) in receptor activation by the *Stachel*. Our study shows that these residues not only regulate receptor activation by P-15 but also control receptor surface expression and degradation.

## Results

### Mutations in ECL2 alter the expression of ADGRG2

We transiently expressed the ADGRG2-P622 (WT) and its ECL2 mutants, CWA, CAI, and CAA in HEK cells and used a cell-based ELISA assay to determine their surface expression via their N-terminal HA-tag. We noticed that all ECL2-mutants show significantly reduced expression on the surface in comparison to the WT receptor (Figure 2a). This lower surface expression was confirmed by immunofluorescence imaging of receptors on the cell surface (Figure 2b). Measuring the total expression via the V5-tag at the C-terminus by ELISA revealed that all three mutants are expressed at reduced levels compared to the WT receptor (Figure 2c). Western blotting of the whole-cell lysates confirmed the reduced expression of mutants (Figure 2d). Together, these data suggest that the expression of ADGRG2 is controlled by tryptophan and isoleucine in the ECL2.

**Figure 2.**
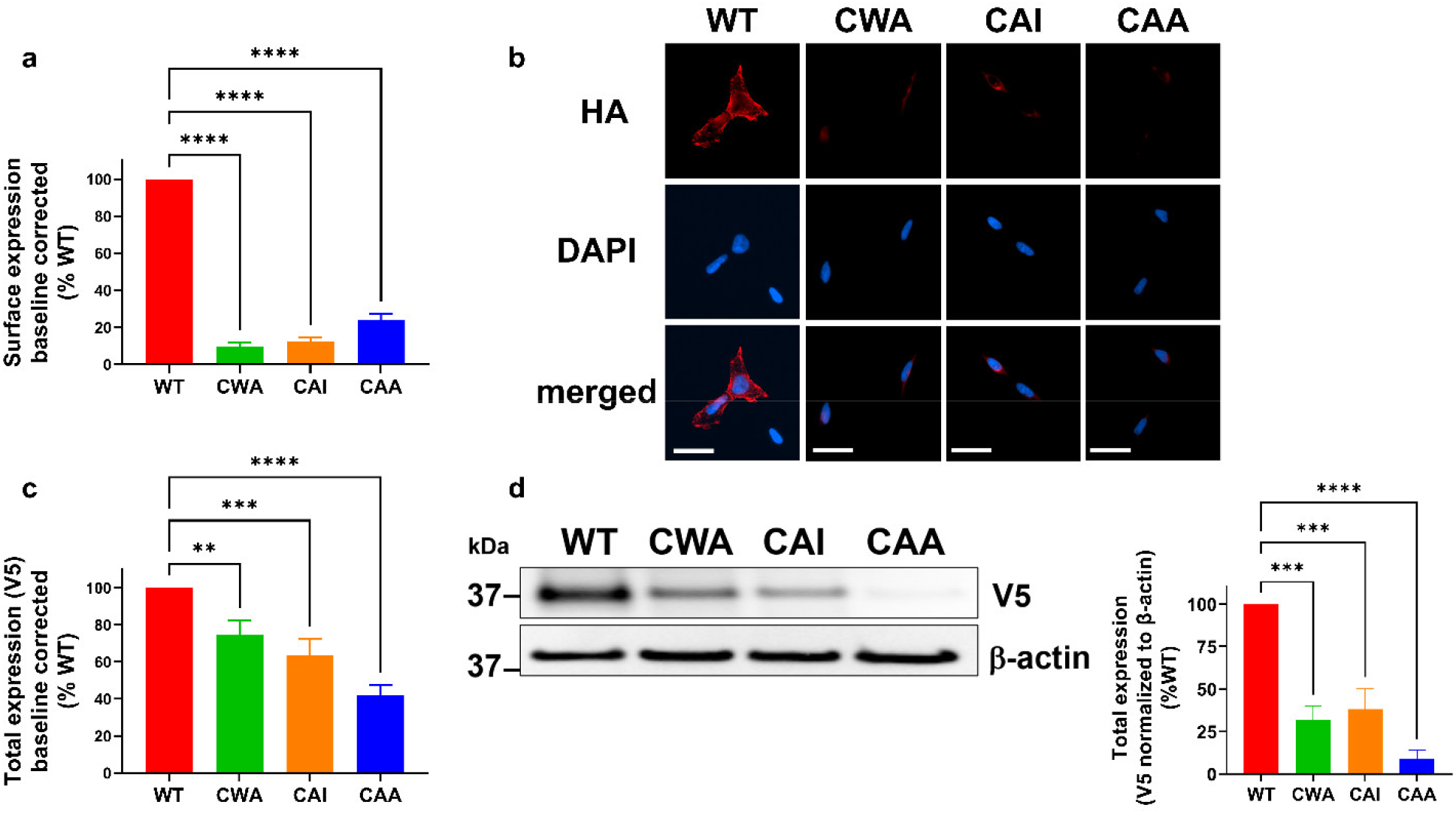
Expression of ADGRG2-P622 is regulated by the residues in the CWI motif in the ECL2. HEK cells were transfected with the same dose of plasmids expressing WT or mutant ADGRG2-P622. Cell surface expression of receptors was determined by (a) ELISA and (b) immunofluorescence imaging using an antibody against the N-terminal HA-tag in non-permeabilized conditions (scale bars: 20 μm). (c) ELISA was used to measure the total expression by using an antibody against the C-terminal V5 tag in permeabilized condition. (d) Representative blots show C-terminal V5-tagged WT and mutant receptors in total cell lysates. Densitometry data were used to normalize V5 expression to β-actin and then to WT plasmid. For ELISA assays, data are corrected for baseline (empty plasmid) values and then normalized to that of WT plasmid. For bar graphs, data are mean ± S.E.M from 4 (for a and c, performed in quadruplicate) and 3 (b and d) independent experiments. One-Way ANOVA with Dunnett’s multiple comparison test was used for statistical analyses. *:P<0.05; **:P<0.01; ***:P<0.001; ****:P<0.0001.

### ECL2-mutations accelerate ADGRG2 degradation

To determine if the reduced expression is due to possible degradation of ECL2 mutants, we isolated cell lysates at several time points post-incubation with a protein translation inhibitor, cycloheximide. The expression of the WT receptor remained substantially unchanged up to 8 hours post-inhibition of translation. However, the expression of all ECL2 mutants was significantly reduced starting at 2 hours post-inhibition and continued to diminish continuously up to 8 hours (Figure 3). This set of data reveals that mutation of conserved residues in the ECL2 of ADGRG2 increases receptor degradation.

**Figure 3.**
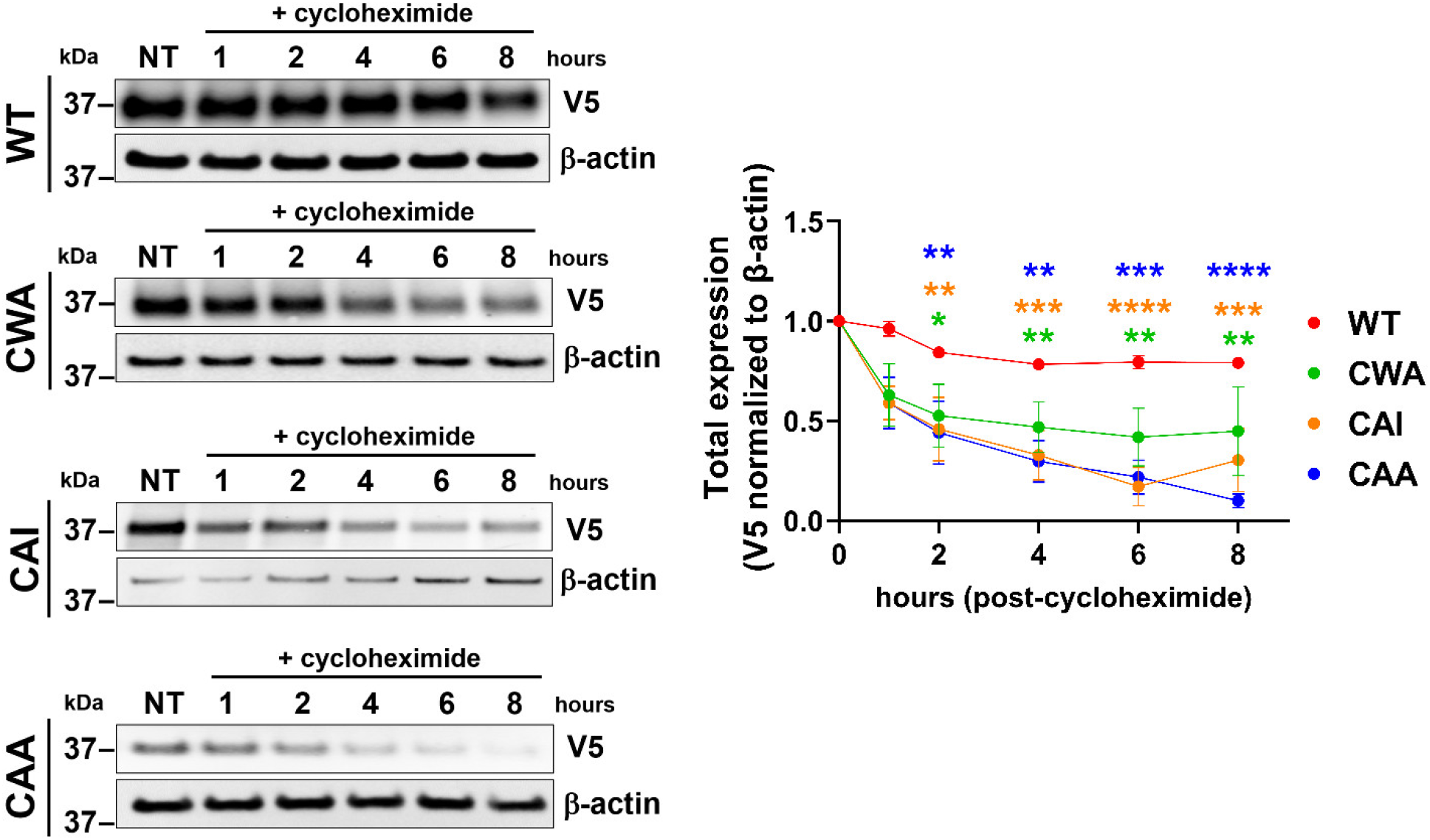
Degradation of ADGRG2-P622 is regulated by the residues in the CWI motif in the ECL2. HEK cells were transfected with the same dose of plasmids expressing WT or mutant ADGRG2-P622. Cells were then incubated with cycloheximide (100 μg/ml) for up to 8 hours and total cell lysates were run on SDS-PAGE, transferred to PVDF membranes, and probed with V5 and b-actin antibodies. Representative blots show reduction of mutant receptor levels over time. Densitometry analyses are shown on the right. Data are mean ± S.E.M from 4 independent experiments. Two-Way ANOVA with Dunnett’s multiple test was used to compare expression levels at each time point to that of basal expression (NT) of respective receptors. *:P<0.05; **:P<0.01; ***:P<0.001; ****:P<0.0001.

### Mutation of tryptophan and isoleucine in ECL2 ablates ADGRG2-mediated induction of transcription factors

We have previously shown that the synthetic P-15 induces cAMP-response element binding protein (CRE) transcription factor in a luciferase reporter assay. To test whether ECL2 mutations affect receptor activation by the *Stachel* peptide despite their reduced surface expression, we used two approaches. First, we normalized the CRE induction downstream of P-15-activated WT and CAA mutants to that of their surface expression (Supplementary Figure 2). This approach revealed a lack of activation of CAA mutant by P-15. Second, we adjusted the dose of the transfected plasmids so that the mutants and WT receptors reach similar surface expression levels (Figure 4a). Thereafter, for all signaling assays, we adjusted the doses of mutant and WT plasmids and supplemented the total plasmid dose with empty backbone pcDNA3.1 plasmid.

**Figure 4.**
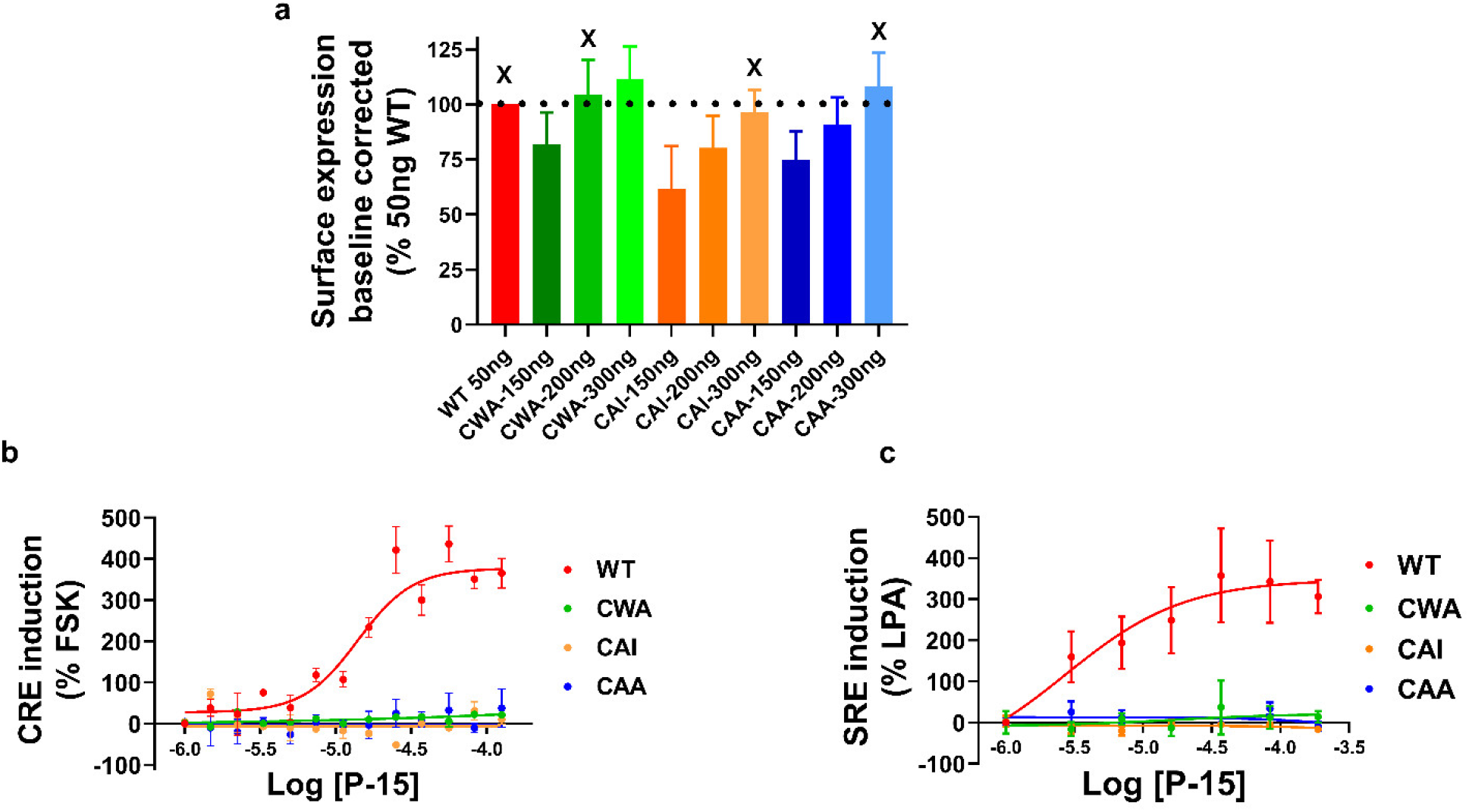
ECL2 plays a major role in ADGRG2 activation by *Stachel* peptide. (a) HEK cells were transfected with one dose of WT and different doses of mutant ADGRG2-P622 plasmids. Cell surface expression of receptors was determined by ELISA using an antibody against the N-terminal HA-tag in non-permeabilized conditions. Data are corrected for baseline (empty plasmid) values and then normalized to that of 50 ng WT-transfected cells. X denotes the amount of mutant plasmids that resulted in comparable surface expression as 50 ng WT plasmid. Data are mean ± S.E.M from 3 independent experiments performed in quadruplicate. (b & c) Cells were transfected with adjusted doses of either WT or mutant plasmids along with either CRE-Luc (b) or SRE-Luc (c) plasmids. After an overnight of serum starvation, cells were activated with increasing concentrations of P-15 for 5 hours. Luciferase induction was measured in a luminescence-based assay. Relative light units recorded in a luminometer were normalized to that of either 10 μM forskolin (FSK) or 10 μM lysophosphatidic acid (LPA) and are presented as mean ± S.E.M from 3 independent experiments performed in triplicate.

Using the CRE and SRE reporter assays, we stimulated cells expressing comparable surface levels of WT and mutant receptors with increasing concentrations of P-15 or vehicle for 5 hours. The basal CRE and SRE activity was similar among WT and mutant receptors. Concentration-response curves revealed that while the WT receptor responds in a concentration-dependent manner to P-15, none of the mutants are activated by this synthetic peptide (Figure 4b and c). These data suggest that activation of some pathways downstream of ADGRG2 is regulated by the conserved residues in the ECL2.

### Gαs pathway activation by the Stachel is controlled by the ECL2

We have previously used an endpoint homogenous time-resolved FRET (HTRF) assay to measure cAMP production downstream of ADGRG2-P622-Gαs pathway^11^. To monitor the production of cAMP in kinetic mode, we developed a new cAMP production assay by using a genetically-encoded biosensor^20^. We induced cAMP production by forskolin, an activator of adenylyl cyclase, and optimized the amount of sensor needed to reach a robust adenylyl cyclase response (Supplementary Figure 3). A 20-second baseline measurement was followed by stimulation with increasing concentrations of P-15 in cells that were transfected with adjusted doses of WT and ECL2-mutated receptors (Figure 5a-d). While the WT receptor responded to P-15 in a concentration-dependent manner, the mutants showed no response, except for the CWA mutant that responded marginally to the highest P-15 concentration. To ensure that increased doses of transfected plasmids did not alter the responsiveness of the adenylyl cyclase-cAMP axis, we compared the forskolin-induced cAMP production. Our data show that despite increased doses of transfected mutant plasmids, cells respond similarly to the forskolin (Figure 5e).

**Figure 5.**
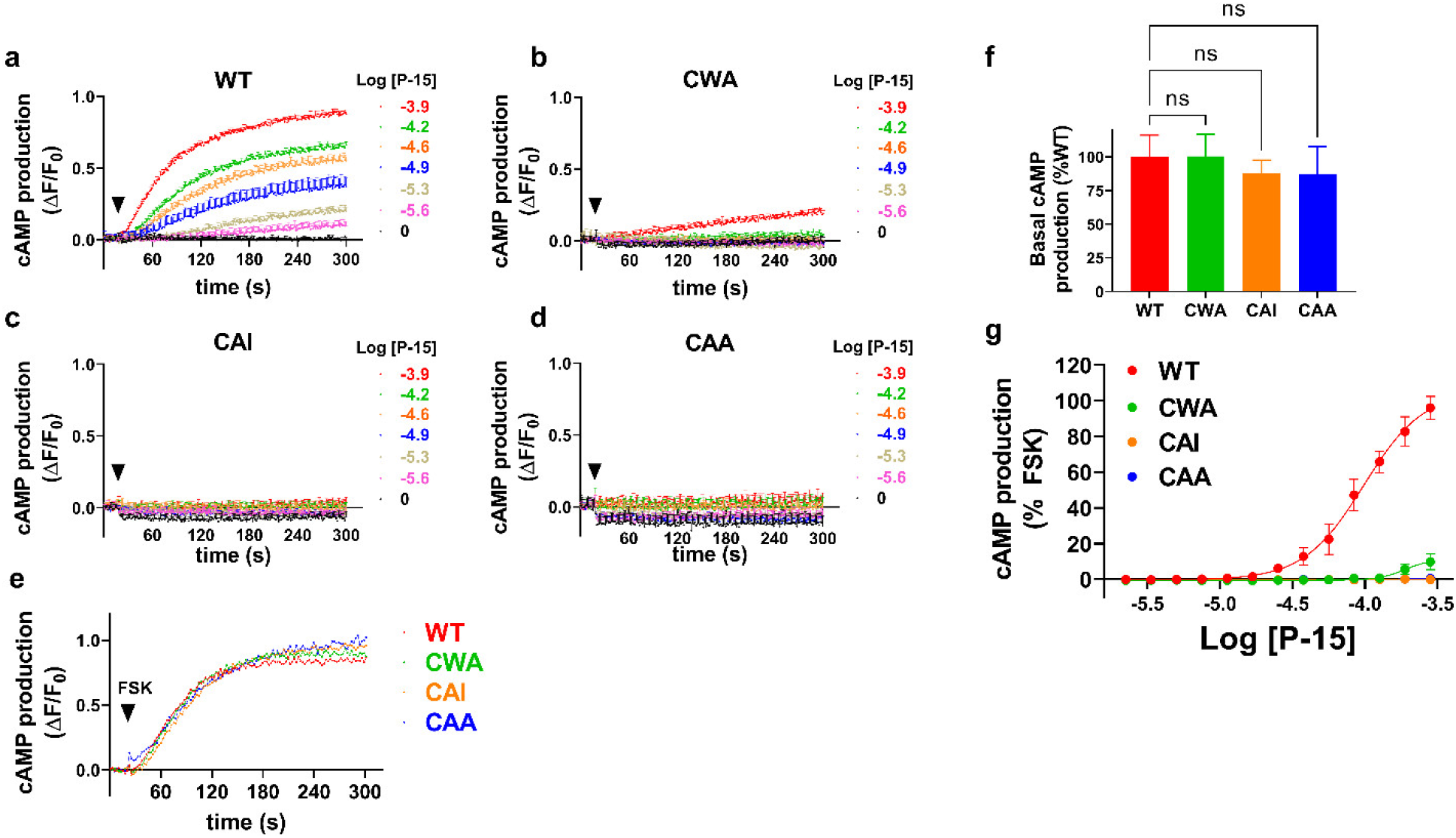
ECL2-mutated ADGRG2-P622 receptors are not activated by P-15. (a-d) Cells were transfected with adjusted doses of WT and mutant plasmids and were transduced with a cAMP biosensor overnight. After a 20-second basal recording of fluorescence (Ex: 488nm; Em: 525nm), cells were activated with increasing concentrations of P-15 (arrowheads), and fluorescence was recorded for another 280 seconds. Relative fluorescence unit (RFU) data were analyzed in GraphPad Prism and are presented as change in RFU divided by the initial RFU (ΔF/F0). (e) Cells transfected with adjusted doses of plasmids and biosensor were stimulated with 1 μM forskolin (FSK). Data are mean ± S.E.M and are representative of 3 independent experiments performed in triplicate. (f) Basal cAMP production was measured after overnight incubation with 0.5 mM IBMX in starvation media in an HTRF assay. Relative cAMP production is presented as mean ± S.E.M from three individual experiments performed in triplicate. Data were compared with WT with one-Way ANOVA with Dunnett’s test. ns: not significant. (g) Cells were stimulated with increasing concentrations of P-15 for 2 hours and cAMP production was measured in an HTRF assay. Data were normalized to the response induced by 1 μM forskolin (FSK) and are mean ± S.E.M from three independent experiments performed in duplicate.

Using the HTRF assay in the presence of IBMX, an inhibitor of phosphodiesterases, to accumulate cAMP production overnight, we found that the basal levels of cAMP are comparable among receptors (Figure 5f). A 2-hour stimulation with P-15 results in a concentration-dependent production of cAMP by the WT receptor (Figure 5g) and a modest response by CWA mutant. Together, these data show that induction of the Gαs-adenylyl cyclase-cAMP pathway by *Stachel* is dependent on the presence of an intact ECL2.

### ECL2 mutations derail the whole-cell response to P-15

We and others have shown that ADGRG2 couples to multiple signaling pathways including Gαq, Gαs, and Gα13^7,11,21,22^. Considering that measuring the production of either a single second messenger (e.g. cAMP) or induction of a couple of transcription factors (e.g. CRE and SRE) cannot detect the ‘overall’ response of cells to GPCR ligands, whole-cell label-free assays have been developed^23–26^. To investigate if the overall response of cells to synthetic P-15 is regulated by the ECL2 conserved residues, we developed a new non-invasive whole-cell label-free impedance assay. Activation of WT-transfected cells with P-15 resulted in a rapid decrease in cell monolayer impedance to 1kHZ frequency of electrical field voltage (Figure 6). However, neither empty plasmid nor ECL2-mutated receptors responded to the P-15 stimulation. This data confirms the lack of receptor activation by *Stachel* peptide in the absence of ECL2 conserved tryptophan and isoleucine residues.

**Figure 6.**
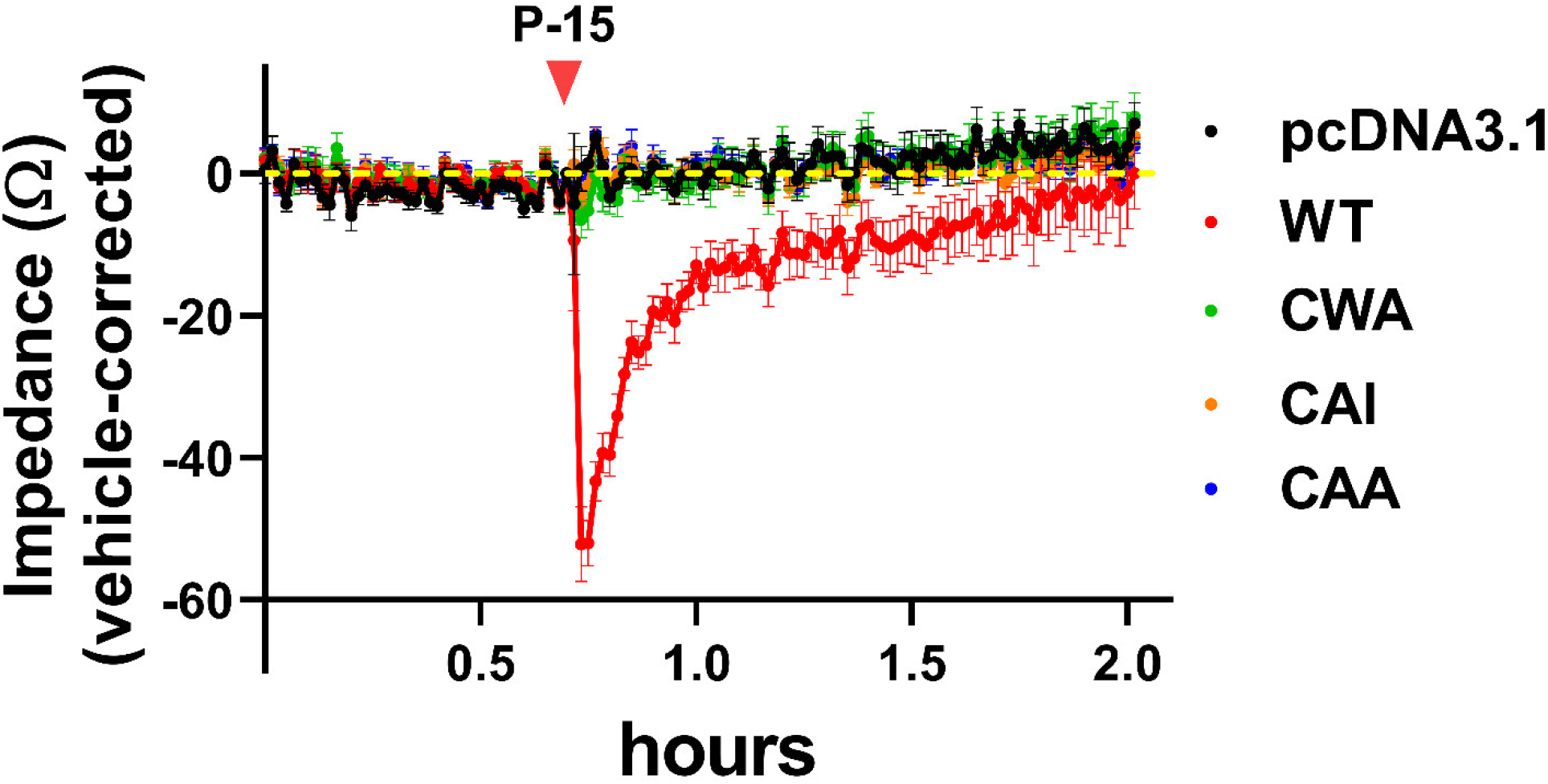
Whole-cell impedance assay shows a lack of activation of the ECL2-mutated ADGRG2 receptor by the *Stachel* peptide. Cells were transfected with adjusted doses of WT and mutant plasmids and were then seeded in plates that have electrodes at the bottom of each well to apply various frequencies of electric voltages. Serum-starved cells, at monolayer confluency, were kept in Maestro Z device until the recorded impedance (Ω) reached a steady state. Cells were stimulated with either vehicle (DMSO) or 100 μM P-15 and impedance was recorded for up to 2 hours. Data are corrected to that of the vehicle for each plasmid and are mean ± S.E.M from a representative experiment out of 3 independent experiments conducted in quadruplicate.

## Discussion

Our understanding of the mechanisms of activation of aGPCRs has substantially increased over the last decade. This progress was made possible due to basic pharmacology and cell biology studies in various cellular backgrounds and model organisms^8,9,27–29^. Now we know that binding of an extracellular molecular partner to the NTF of ADGRG1 (GPR56) dissociates it from the CTF and unmasks a *Stachel* sequence, which subsequently activates ADGRG1^14,15,30^. Similar mechanisms of activation exist for a few other aGPCRs^7,31^. However, the binding site of the *Stachel* on the cognate aGPCR remains largely unknown. Structural similarities of class B1 GPCRs to aGPCRs (formerly class B2) and the fact that class B1 receptors bind to peptide hormones provides hints on the possible binding site of the *Stachel*. Specifically, the ECL2 forms an orthosteric binding pocket for peptide hormones in PTH 1 receptor (PTH1R)^32^ and Corticotropin-releasing factor 1 receptor (CRFR1)^33^. In the current study, we tested the role of conserved residues in and around the CW motif of the ECL2 in the activation of an NTF/*Stachel*-truncated ADGRG2 (ADGRG2-P622) by its *Stachel*, P-15. Our results provide evidence that the lack of isoleucine and/or tryptophan in the ECL2 not only abolishes the signaling pathways of ADGRG2 but also dysregulates receptor stability in HEK293 cells.

We chose the ADGRG2-P622 receptor for this study because we had previously shown that ADGRG2-P622 responds significantly larger to the synthetic P-15 in comparison to the full-length (ADGRG2-FL) or truncated ADGRG2-ΔNTF in a battery of assays^11^. This maximal response by ADGRG2-P622 is presumably due to the availability of the binding site for P-15, which might be either masked by the NTF in ADGRG2-FL or occupied by the endogenous P-15 in ADGRG2-ΔNTF. The lack of constitutive activity in cells that express ADGRG2-P622 in the current study confirms the role of endogenous *Stachel* in the activation of ADGRG2.

Here, we used several readouts to assess receptor response to the synthetic peptide. The use of kinetic and endpoint cAMP production assays allowed us to measure the Gαs signaling of ADGRG2. Genetically encoded biosensor and HTRF assays showed a lack of activation of ECL2 mutants either in the short-term (minutes) or long-term (2 hours). We supplemented these assays with CRE and SRE reporter assays, in which cells are activated for 5 hours. Both assays showed a lack of ECL2 activation by P-15, suggesting that the coupling of both Gαs and Gα13 proteins to ADGRG2 is ablated in the absence of either tryptophan or isoleucine or both. To assess other possible but unknown signaling pathways that may originate from ADGRG2, we developed a whole-cell label-free impedance assay. We have previously used similar label-free assays to study GPR55 and cannabinoid 2 receptor^34,35^. This method allowed us to measure the overall response, rather than a singular signaling pathway, from a monolayer of cells upon activation with P-15 in kinetic mode. Using this readout, we observed a strong fast *Stachel*-specific reduction of impedance from WT receptor-expressing cells, indicating possible movement of cells and consequent disruption of cell-cell contact in the monolayer. This response was transient, and impedance was mostly restored in 15 min. Nevertheless, we did not detect a response from cells expressing any of the ECL2 mutated receptors, further confirming the lack of activation by P-15.

It is noteworthy that mutant receptors were degraded in the absence of either endogenous or synthetic *Stachel* peptide. Do the ECL2 mutations induce constitutive activity followed by sustained internalization and degradation? This does not seem to be plausible because we did not detect any altered basal activity either in CRE reporter assay or cAMP production assays. Do the ECL2 mutations hinder receptor transport and incorporation into the plasma membrane? Our western blotting, ELISA, and immunofluorescence imaging data suggest that the diminished surface expression of mutants is likely to be due to the fast degradation of receptors upon translation. We have previously shown that truncated ADGRG2-ΔNTF gets ubiquitinated but is expressed on the plasma membrane at levels comparable to that of the full-length ADGRG2^11,12^. Further studies are required to unravel 1) the mechanisms by which ADGRG2 gets degraded (lysosomal or proteasomal^36^) and 2) how the isoleucine and tryptophan of ECL2 impact these mechanisms. Irrespective of the underlying reason for diminished total expression and degradation, we adjusted the surface expression levels by increasing the dose of the transfected plasmids and conducted our signaling assays in this condition. Nevertheless, comparable surface expression of ECL2-mutated and WT receptors did not rescue their response to P-15, supporting the role of ECL2 in *Stachel*-mediated activation of ADGRG2. Whether the ECL2 of other aGPCRs play similar roles in receptor stability and activation by *Stachel* peptides warrants further studies.

While finalizing this manuscript, we noticed that a separate group reported that several residues, including the tryptophan and isoleucine of the ECL2, participate in the binding of *Stachel* peptide to ADGRG2^37^. Although this study corroborates some of our findings, differences exist between the two studies. We used ‘human’ ADGRG2-P622 that lacks the NTF and the *Stachel* sequence to prevent interaction of the endogenous P-15 on the N-terminus of the receptor with the receptor. *Sun et. al.*, however, determined the binding of a ‘mouse’ P-15 that was ‘modified’ at several residues to the ‘mouse’ ADGRG2 ortholog. This modified P-15 showed higher potency compared to natural P-15 and is potentially a new pharmacological tool for future ADGRG2 studies. However, its application remains to be validated by testing whether it shows affinity or activity towards other aGPCRs that share similar *Stachel* sequence and ECL2 motifs as ADGRG2. Another major difference between these two studies is that while we observed a significant alteration of ECL2-mutated receptor stability and cell surface expression, *Sun et. al.* did not report such changes. These differences make a direct comparison of the two studies challenging.

ADGRG2 plays various physiological and pathological roles. For instance, mutations or truncations in ADGRG2 are associated with different forms of male infertility^38,39^ and male *Adgrg2*-null mice are infertile^40^. ADGRG2 also promotes proliferation and metastasis of Ewing sarcoma cells both *in vitro* and *in vivo*^41^. We have previously shown that ADGRG2 is enriched in parathyroid glands, is upregulated in parathyroid adenomas from patients with primary hyperparathyroidism, and its activation by P-15 elevates PTH secretion^12^. Although ADGRG2 is still an orphan aGPCR, our findings on the role of ECL2 in receptor stability and *Stachel*-mediated activation provide a foundation for future targeting of this receptor via small molecules or biologics.

## Methods

### Cell culture and transfection

AD-293 (HEK) cells were purchased from Agilent Technologies (Santa Clara, CA, USA #240085) and cultured in DMEM media (Sigma, St. Louis, MO, USA #D6429) supplemented with 10% FBS, 100 U/mL penicillin, and 100μg/mL streptomycin (Thermo Fisher Scientific, Waltham, MA, USA #15140-122). Cells were transfected with plasmids using lipofectamine 2000 reagent (Thermo Fisher Scientific #11668019) following the manufacturer’s instructions.

### Antibodies

Antibodies were purchased from the following sources: Cell Signaling Technologies (Beverly, MA, USA): rabbit anti-HA (#3724); Thermo Fisher Scientific: mouse anti-V5 (#R960-25); Sigma: rabbit anti-β-actin (#A2066); Biolegend (San Diego, CA, USA): mouse anti-HA (#901513).

### Receptor mutagenesis

We used the Q5 site-directed mutagenesis kit (NEB, Ipswich, MA #E0552S) to generate mutations in the ECL2 of human ADGRG2. Our template for mutations was pcDNA3.1-3xHA-P622-V5 (WT)^11^, a plasmid that expresses an NTF/*Stachel*-truncated ADGRG2 with an N-terminal 3xHA tag and a C-terminal V5 tag. The following primer pairs were used to construct mutants: P622-CWA^800^ (*for*: CTTCTGCTGGGCCAACAACAATGC; *rev*: TCATCCGGTGAACCATTG), P622-CA^779^I (*for*: TGACTTCTGCGCCATCAACAACAATGCAGTATTC; *rev*: TCCGGTGAACCATTGGGG), P622-CA^779^A^800^ (*for*: TGACTTCTGCGCCGCCAACAACAATGCAGTATTCTAC; *rev*: TCCGGTGAACCATTGGGG). The resulting CWA, CAI, and CAA constructs were verified by sequencing both strands.

### On-Cell ELISA

Cells were seeded in 96 well plates and transfected with 100 ng of one of the following plasmids: pcDNA3.1 (empty vector), WT, CWA, CAI, and CAA. Twenty-four hours after transfections, cells were starved using DMEM (Thermo Fisher Scientific #21068028) supplemented with glutamine and 1.25mM Ca^2+^ overnight. Cells were then fixed with 4% paraformaldehyde for 15 minutes at room temperature. After several washes with TBS, cells were blocked for 30 minutes in TBSM (TBS + 3% milk) for surface staining or TBSM supplemented with 0.2% Triton X-100 for total staining. Cells were then incubated with either rabbit anti-HA (1:2000) or mouse anti-V5 (1:2000) antibodies in TBSB (TBS + 3% BSA) for 2 hours at room temperature. After several washes, cells were incubated for 1 hour with 1:2000 dilution of either horseradish peroxidase (HRP)-linked horse anti-mouse IgG (Cell Signaling Technologies #7076) or HRP-linked goat anti-rabbit IgG (Cell Signaling Technologies #7074) antibodies in TBSM. After 5 washes with TBS, cells were incubated with 3,3’,5,5’-Tetramethylbenzidine (TMB) (Sigma #t0440) for 5 min at room temperature. The reaction was stopped by an equal volume of 1 N HCl. Absorbance at 450nm was measured using a FlexStation III plate reader. In a separate set of experiments, cells were transfected with several amounts of the aforementioned plasmids to determine the doses of mutant plasmids that result in similar surface expression compared to WT.

### Degradation assay

Cells were transiently transfected with plasmids in 6-well plates and 24 hours post-starvation were incubated with 100μg/ml of cycloheximide (Sigma #4859) for indicated time points. Cells were lysed using radioimmunoprecipitation assay (RIPA) lysis buffer (EMD Millipore, Billerica, MA, USA #20-188) supplemented with cocktails of protease (Thermo Fisher Scientific #78429) and phosphatase inhibitors (EMD Millipore #524625). Lysates were then centrifuged at 13,000 rpm for 5 min at 4°C and supernatants were collected.

### Western blotting

Protein lysates were boiled with reducing sample buffer (Thermo Fisher Scientific #NP0007 and #NP0009) for 10 minutes and loaded on 4-12% Bis-Tris gels (Thermo Fisher Scientific #NP0336BOX) then transferred to polyvinylidene fluoride (PVDF) membranes. Blocking was performed in TBSTM (TBS + 0.1% Tween-20 + 10% nonfat dry milk) followed by incubation with primary antibodies in TBSTM (1:2000 for HA and V5; 1:500 for β-actin), overnight at 4°C. To detect primary antibodies, cells were incubated with HRP-linked horse anti-mouse IgG and HRP-linked goat anti-rabbit IgG antibodies (1:5,000) in TBSTM for 2 hours at room temperature. Blots were washed and then developed with ECL SuperSignal™ West Femto substrate (Thermo Fisher Scientific #34095). Blots were imaged and analyzed in iBright™ FL1500 Imaging System (Thermo Fisher Scientific).

### Immunofluorescence staining and imaging

HEK cells were transfected with plasmids, seeded on glass coverslips, and serum-starved overnight. Cells were fixed in 4% paraformaldehyde for 10 min and were washed with PBS several times. Cells were blocked with 5% goat serum in PBS for 1 hour and incubated with mouse anti-HA antibody (1:1000) in 1% BSA in PBS for 2 hours at room temperature. After several washing steps with PBS, HA antibody-bound receptors were labeled with Alexa Fluor 594–conjugated goat anti-mouse antibody (1:500) in 1% BSA in PBS for 1 hour. Cells were mounted in ProLong Diamond Antifade Mountant with DAPI (ThermoFisher Scientific #P36971) for nuclear counterstaining. Fluorescence microscopy was conducted by 40x oil objective (1.4 NA) on a Nikon Ti-E microscope equipped with a 16.2 MegaPixels DS-Ri2 camera and images were analyzed with Nikon NIS-Elements Basic Research software.

### Luciferase reporter assays

Luciferase reporter assay was performed as described previously^12^, with some modifications. HEK cells were seeded in white clear-bottom 96-well plates (20,000 cells/well) and were transfected with adjusted amounts of receptor plasmids along with 100 ng of either pCRE-Luc or pSRE-Luc reporter plasmids. Cells were then stimulated with increasing concentrations of P-15, forskolin, or vehicle for 5 hours at 37°C. Using Steadylite reagents (PerkinElmer, Hopkinton, MA, USA, #6066756), the luminescence was measured in a FLEXStation III device.

### cAMP production assays

Cells were resuspended in DMEM + 10% FBS and were mixed with Upward Green cADDis cAMP Assay Kit BacMam sensor (Montana Molecular, Bozeman, MT, USA #U0200G), supplemented with 2mM sodium butyrate. Cells were then seeded at 50,000 cells/well in black, clear-bottom, 96-well plates (150μl/well), incubated for 30 min at room temperature in the dark, and then at 37°C for 24 hours. Cells were washed with assay buffer (HBSS; Thermo Fisher Scientific #14065056 supplemented with 20mM HEPES) and kept in dark to acclimate to room temperature for 1 hour in 100 μl/well of assay buffer. Initial assay development was conducted by transducing with various amounts of the BacMam Gαs sensor followed by stimulation with 10 μM forskolin for 800 seconds (Supplementary Figure 3). Considering the speed and amplitude of the signal, we chose 15 μl per well of the sensor as the appropriate volume to monitor cAMP production in HEK cells. After transfection with plasmids, cells were transduced with 15 μl of the sensor (as described above) and were stimulated with increasing concentrations of P-15 the next day. cAMP production was recorded for 280 seconds after an initial 20 seconds of basal measurement of fluorescence at Excitation 488nm and Emission 525nm using the FLEXStation III plate reader. In a different set of experiments, a previously described homogenous time-resolved FRET (HTRF) assay^11^ was used to measure cAMP production either at basal condition (overnight incubation with 0.5 mM IBMX; Sigma #I5879) or after 2 hours of activation with P-15.

### Whole-cell label-free impedance assay

Cells were transfected with adjusted amounts of plasmids and were seeded at a density of 50,000 cells per well in 100 μl of DMEM (Thermo Fisher Scientific #21068028) supplemented with glutamine and 1.25mM Ca^2+^ in CytoView-Z 96-well plates (Axion BioSystems, Atlanta, GA, USA #Z96-IMP-96B) overnight. After 45 minutes of baseline recording of impedance in the Maestro Z device (Axion BioSystems) cells were activated with 100 μM P-15 or vehicle (DMSO). Impedance against 1kHz voltage frequency was recorded for several hours at 37°C, 5% CO2 in a humidified environment in the Maestro Z machine. Data normalization to vehicle and analyses were performed in AxIS Z software.

### Statistical analysis

Statistical analyses were conducted using two-tailed Student’s t-test for comparisons between two groups, One-Way or two-Way ANOVA with Dunnett’s multiple test or Holm-Sidak multiple comparison test in GraphPad Prism 9.0 software; P < 0.05 was considered significant.

## Acknowledgments

We thank the University of Maryland Marlene and Stewart Greenebaum Comprehensive Cancer Center for continuous support. This work was supported by NIH/NIGMS Grant (R01GM130617) to N.B.

## Author contributions

N.B. designed the research; A.A.G., P.A., and N.B. conducted experiments; A.A.G. and N.B. performed data analysis and wrote the manuscript.

