## Supplemental Figures 1-3 for "Conserved residues in the extracellular loop 2 regulate *Stachel*-mediated activation of ADGRG2"

### Supplementary Figures

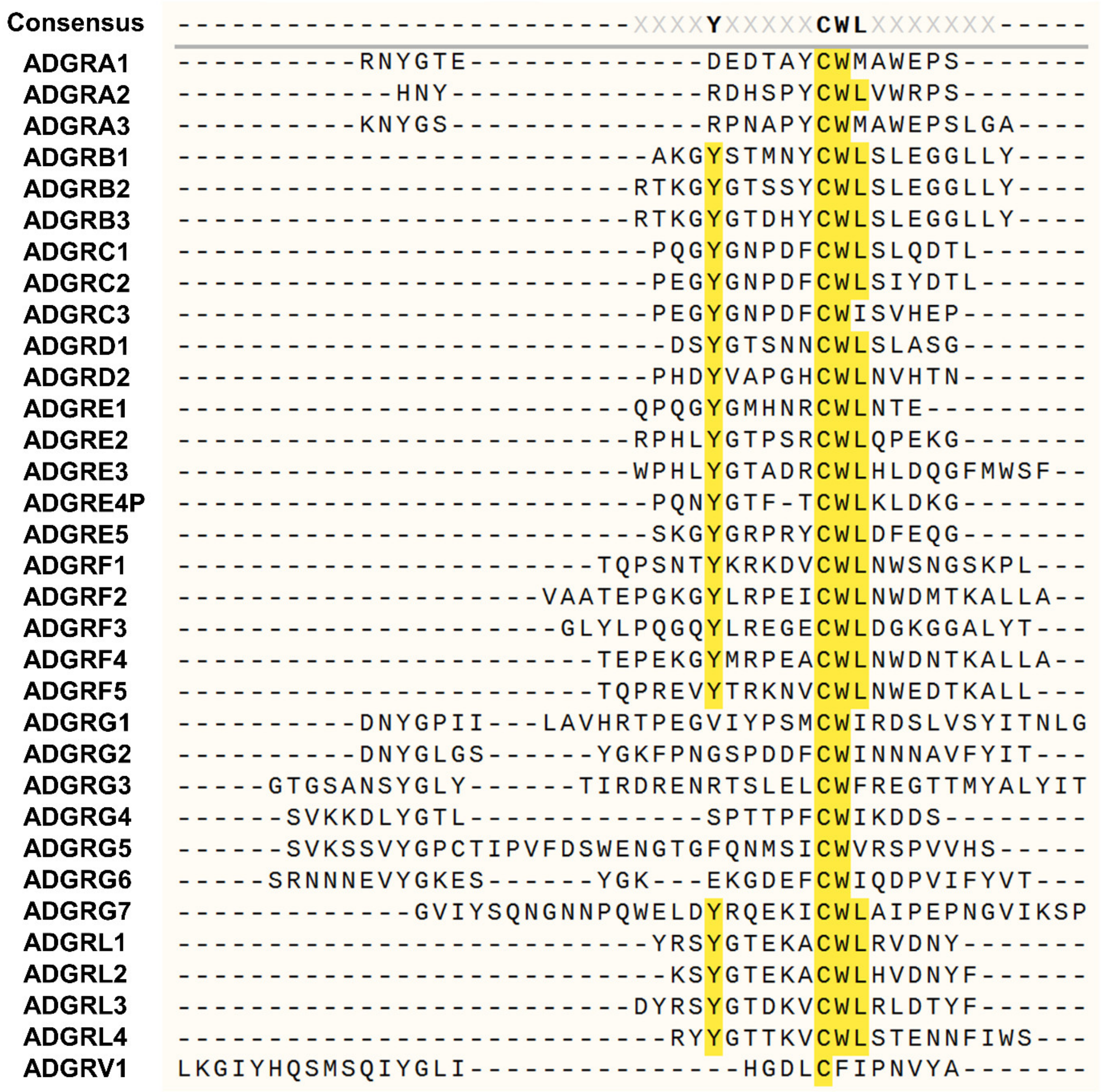


**Supplementary Figure 1.** Multiple alignments of the ECL2 of all 33 members of the aGPCR family. Predicted amino acid sequences of the ECL2 for each aGPCR were derived from Uniprot. The alignment was conducted in SnapGene software using the Clustal Omega algorithm.

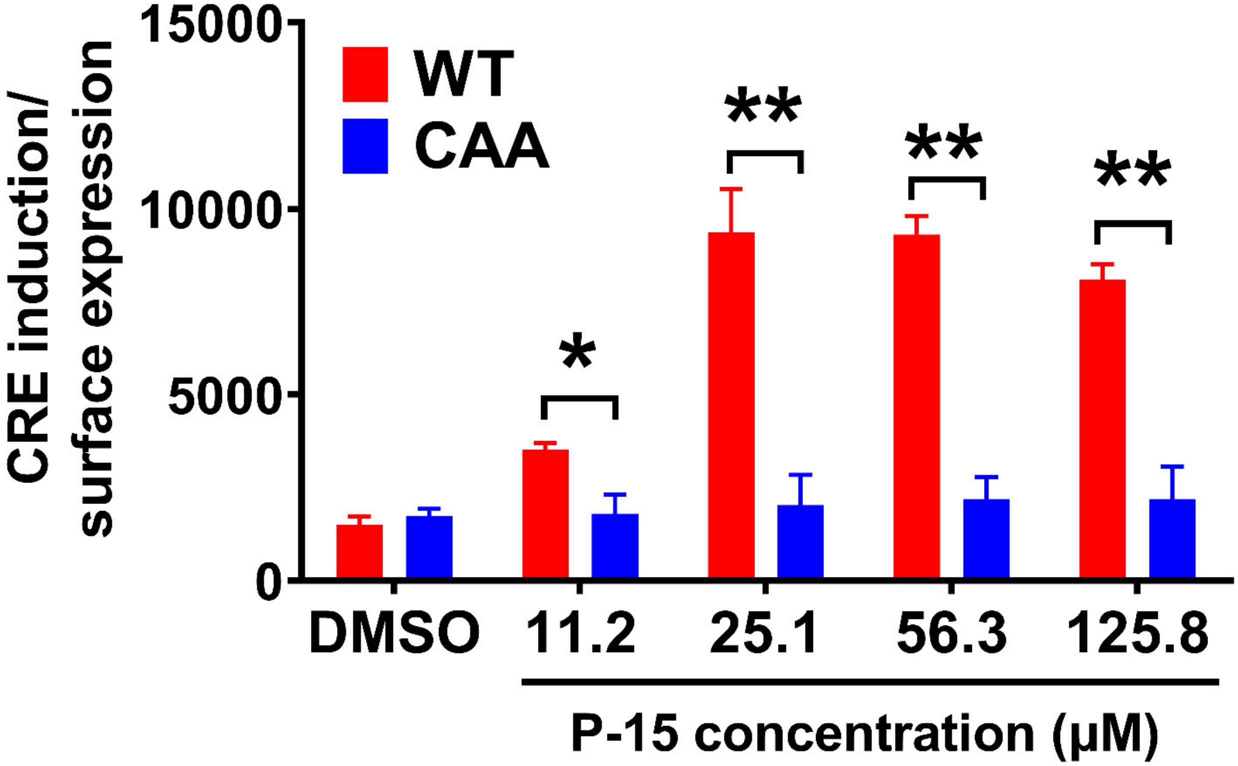


**Supplementary Figure 2.** HEK cells were transfected with the same dose of WT and CAA plasmids along with CRE-Luc plasmid. Luciferase activity in response to increasing concentrations of P-15 was measured in a luminescence-based assay. Relative light units (RLU) were normalized to the surface expression data derived from an ELISA assay and are presented as mean ± S.E.M from 3 independent experiments performed in duplicate. $*P<0.05$, $**P<0.01$. Data were compared with WT at each concentration of P-15 with Holm-Sidak multiple comparison test.

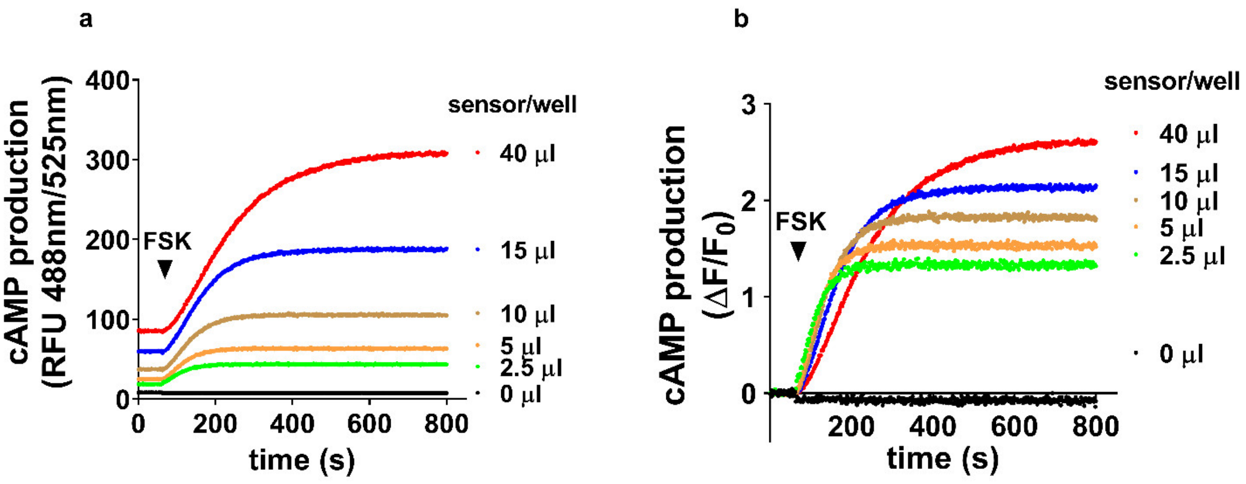


**Supplementary Figure 3.** Characterization of a fluorescent genetically encoded cAMP sensor. HEK cells were transduced with various amounts of Upward Green cADDis cAMP sensor in 96-well plates overnight. Cells were washed with assay buffer and then the fluorescence (Excitation at 488 nm/Emission at 525 nm) was recorded for 1 min before and 12 min after addition of 10 µM FSK. (a) Raw data showing the relative fluorescence units (RFU) for each amount of sensor. (b) Data were analyzed in GraphPad Prism and are presented as change in RFU divided by the initial RFU (ΔF/F0).
